# A self-cooling self-humidifying mosquito carrier backpack for transporting live adult mosquitoes on foot over long distances under challenging field conditions

**DOI:** 10.1101/2024.04.13.588955

**Authors:** Deogratius R Kavishe, Rogath V Msoffe, Goodluck Z Malika, Katrina A Walsh, Lily M Duggan, Lucia J Tarimo, Fidelma Butler, Emmanuel W. Kaindoa, Halfan S Ngowo, Gerry F Killeen

## Abstract

It is often necessary to use motorized transport to move live mosquitoes from distant field collection points into a central insectary, so that their behavioural and/or physiological phenotypes can be assessed under carefully controlled conditions. However, a survey of heritable insecticide susceptibility traits among wild-caught *Anopheles arabiensis* mosquitoes, collected across an extensive study area composed largely of wilderness in southern Tanzania, necessitated that live mosquitoes were carried on foot over distances up to 25 km per day because most of the area was impassable by car, motorcycle or even bicycle during the rains. A self-cooling, self-humidifying carrier backpack was therefore developed that allows live adult mosquito specimens to be transported across rugged miombo woodland and floodplain terrain throughout the year. This wettable backpack was fabricated from stitched Tanzanian *kitenge* cotton fabric and PVC-coated fiberglass netting that allows easy circulation of air in and out. An outer cover flap made of cotton towelling embedded inside a kitenge envelope overhangs the fiberglass netting upper body of the bag, to protect mosquitoes from direct sunlight, and can be soaked with water to maintain low temperature and high humidity inside. Mean survival of insectary-reared female *Anopheles arabiensis* transported through 9 different mobile camps inside the 509 km^2^ Ifakara-Lupiro-Mang’ula Wildlife Management Area (ILUMA WMA), over up to 143km and 25 days, was indistinguishable from those left in the field insectary over the same period. Although considerable variance was observed between different batches of mosquitoes from the insectary and between individual cups of mosquitoes, the different levels and positions inside the backpack had no influence on survival. Temperature and humidity inside the backpack were maintained at standard insectary condition throughout, despite much more extreme conditions immediately outside. When the backpack was applied to collecting wild *An. arabiensis* and *Anopheles quadriannulatus* across a much larger study area of >4000 km^2^, encompassing the ILUMA WMA, some nearby villages and adjacent parts of Nyerere National Park (NNP), it achieved a mean survival rate of 58.2% [95% confidence interval 47.5 to 68.2]. Encouragingly, no difference in survival was observed between ILUMA WMA and NNP even though transport back from NNP involves much longer distances, sometimes involving lengthy journeys by car or boat. Overall, this mosquito carrier backpack prototype appears to represent a viable and effective method for transporting live wild-caught mosquitoes on foot across otherwise impassable terrain under challenging weather conditions with minimal detrimental impact on their survival.

## INTRODUCTION

Malaria vector populations all across the tropics are now resistant to various insecticides, most notably the pyrethroids used to formulate the long-lasting insecticidal nets (LLINs) that have saved so many lives since they were scaled up over the last two decades (Hemingway et al., 2016; Ranson & Lissenden, 2016; WHO, 2023). Due to prolonged, largely singularly reliance upon this exceptionally useful class of active ingredients for a range of public health, veterinary and agricultural applications (Hemingway et al., 2016; Reid & McKenzie, 2016; Urio et al., 2022), pyrethroid resistance now appears ubiquitous across Africa and is clearly eroding the impacts of LLINs (Accrombessi et al., 2023; Killeen & Sougoufara, 2023; Mosha et al., 2022; Tiono et al., 2018).

Looking ahead, pre-emptive insecticide resistance management (IRM) strategies are clearly needed, not only to prevent or slow the emergence of new resistance traits (Killeen, 2020; Killeen & Sougoufara, 2023), but also alleviate (Mangan et al., 2023) and ideally reverse (Kaduskar et al., 2022; Keiding, 1963; Lynch & Boots, 2016; Thornton et al., 2020; White et al., 2014) the selection pressures that currently sustain high frequencies of existing ones. While it is, in principle, possible to restore pyrethroid susceptibility traits among wild vector populations, through back-selection with astutely deployed combinations of complementary insecticide classes (Kaduskar et al., 2022; Keiding, 1963; Lynch & Boots, 2016; Thornton et al., 2020; White et al., 2014), it remains to be seen whether any malaria vector populations still exist in Africa that retain their normal historical diversity of fully insecticide susceptible wild-type genomes.

In order to find such refuge populations of malaria vectors, with genomes that have not yet been bottlenecked by insecticidal selection pressure, it will probably be necessary to look beyond human-settled areas (Epopa et al., 2020), within the remaining well-conserved wilderness areas that are still scattered across the continent. Within the intact natural ecosystems of Africa’s finest national parks, where pesticide use is negligible and wild animals offer mosquitoes diverse alternatives to humans and livestock, desirable wild-type susceptibility traits may well also be conserved. Even on the fringes of such pristine areas, where less rigorous conservation models like community-based Wildlife Management Areas (WMAs) (Lee, 2018; Mwakaje, 2008) result in mixed land cover, such ecologically diverse patchworks of quite different environmental conditions may undermine the impacts of vector control (Killeen & Reed, 2018) and attenuate the selection pressures that underpin emergence of resistance traits (Mangan et al., 2023).

However, due to their typically limited infrastructure and road access, conducting entomological surveys in the most remote and pristine reaches of conservation areas can be challenging especially, when one needs to not only capture adult mosquitoes but also bring them back alive to a central insectary or laboratory for further investigation. The study reported herein therefore responds to that need, as part of a larger project intended to demonstrate that the *portfolio effects* (Killeen & Reed, 2018) caused by wildlife conservation areas allow refugia population of pyrethroid-susceptible malaria vectors to persist, which might spread back into the nearby human-settled area if astute insecticide combination were deployed to favor their survival (Lynch & Boots, 2016; White et al., 2014). In brief, a mosquito carrier backpack was designed and evaluated as a means to transport live wild-collect mosquitoes on foot across an area spanning an extensive community-managed WMA (Lee, 2018; Mwakaje, 2008; Tourism, 2022), nearby villages and an adjacent national park in southern Tanzania (Figure 1), most of which is often otherwise completely inaccessible for much of the year (Duggan, 2024; Walsh, 2024), so that they could be brought back to a central field insectary which is 3.1 km from the nearest camp and up to 229 km from the farthest camp for captive propagation and further investigation of their heritable insecticide susceptibility phenotypes.

**FIGURE 1.**
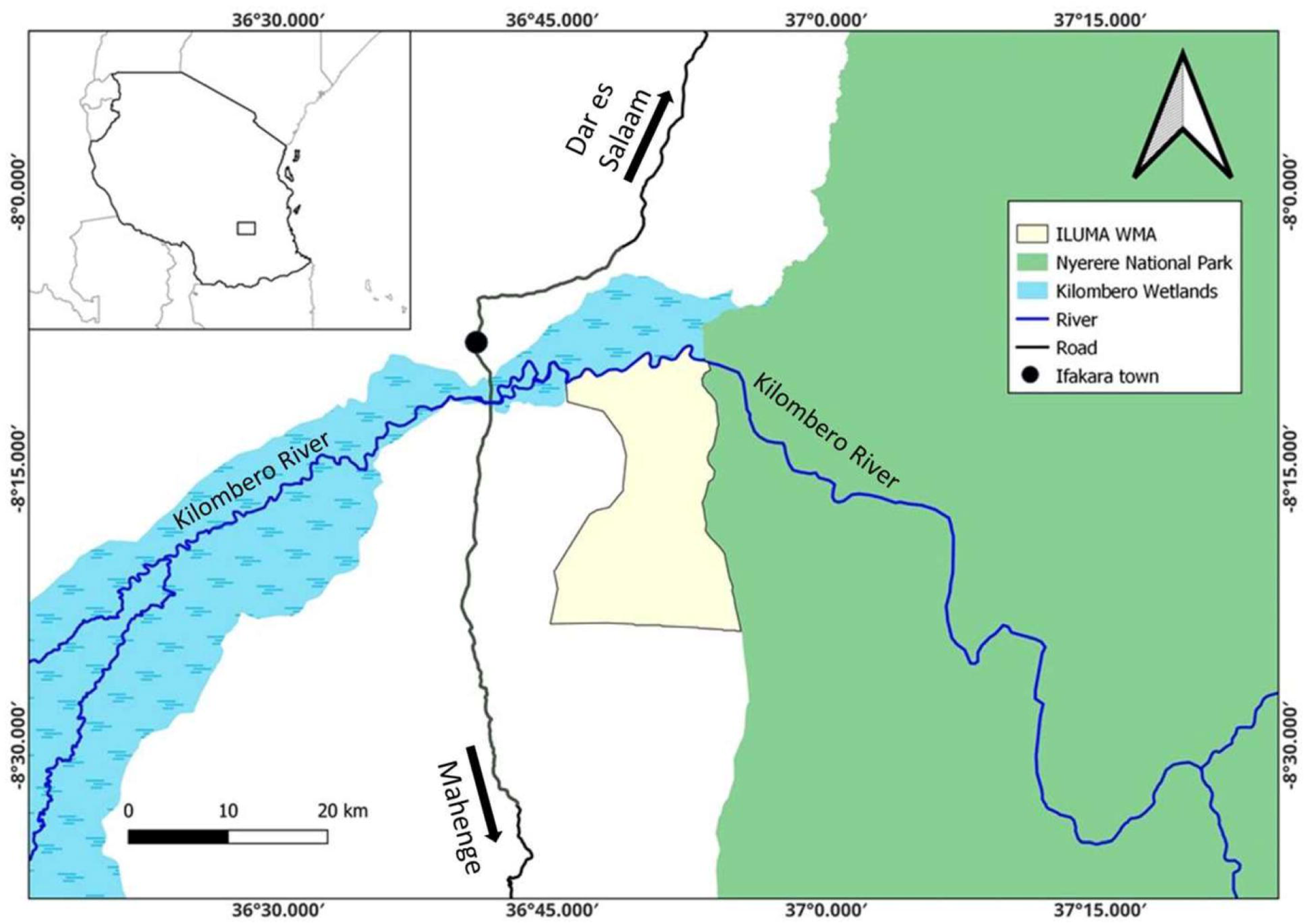
Map displaying the survey area in a national and local context in southern Tanzania. The insert at the top left is a map of Tanzania, while the enlarged area shows the part of the Ifakara-Lupiro-Mang’ula Wildlife Management Area (ILUMA WMA) to the south of the Kilombero River, where most of the surveys were carried out, illustrated in the context of the nearby Nyerere National Park (NNP) to the east, the extensive wetlands of the Kilombero Valley inland delta upstream to the west, and Ifakara town to the north-west.

## MATERIAL AND METHODS

### Mosquito carrier backpack design

The mosquito carrier backpack described herein (Figure 2) was designed using locally available materials, such as cotton fabric known as *kitenge*, cotton towels, polyvinyl chloride (PVC) coated fiberglass netting that is commonly used for window screening, and a one-inch-wide flat metal bar. The metal bar was cut, bent, and welded to make a metal frame with three levels that can hold six paper cups in each level (Figure 2A). To make a complete mosquito carrier backpack the metal frame was placed inside a wettable backpack (figure 2C) tailored from cotton fabric and PVC-coated fiberglass netting that allows easy air circulation in and out. An outer cover flap made of cotton towel embedded inside a *kitenge* fabric envelope hangs over the fiberglass netting on the upper body of the bag to protect mosquitoes from direct sunlight. This absorbent cover flap can also be soaked with water to maintain low temperature and high humidity inside the bag. For these experiments, two mosquito carrier backpacks were made and used to transport mosquitoes.

**FIGURE 2.**
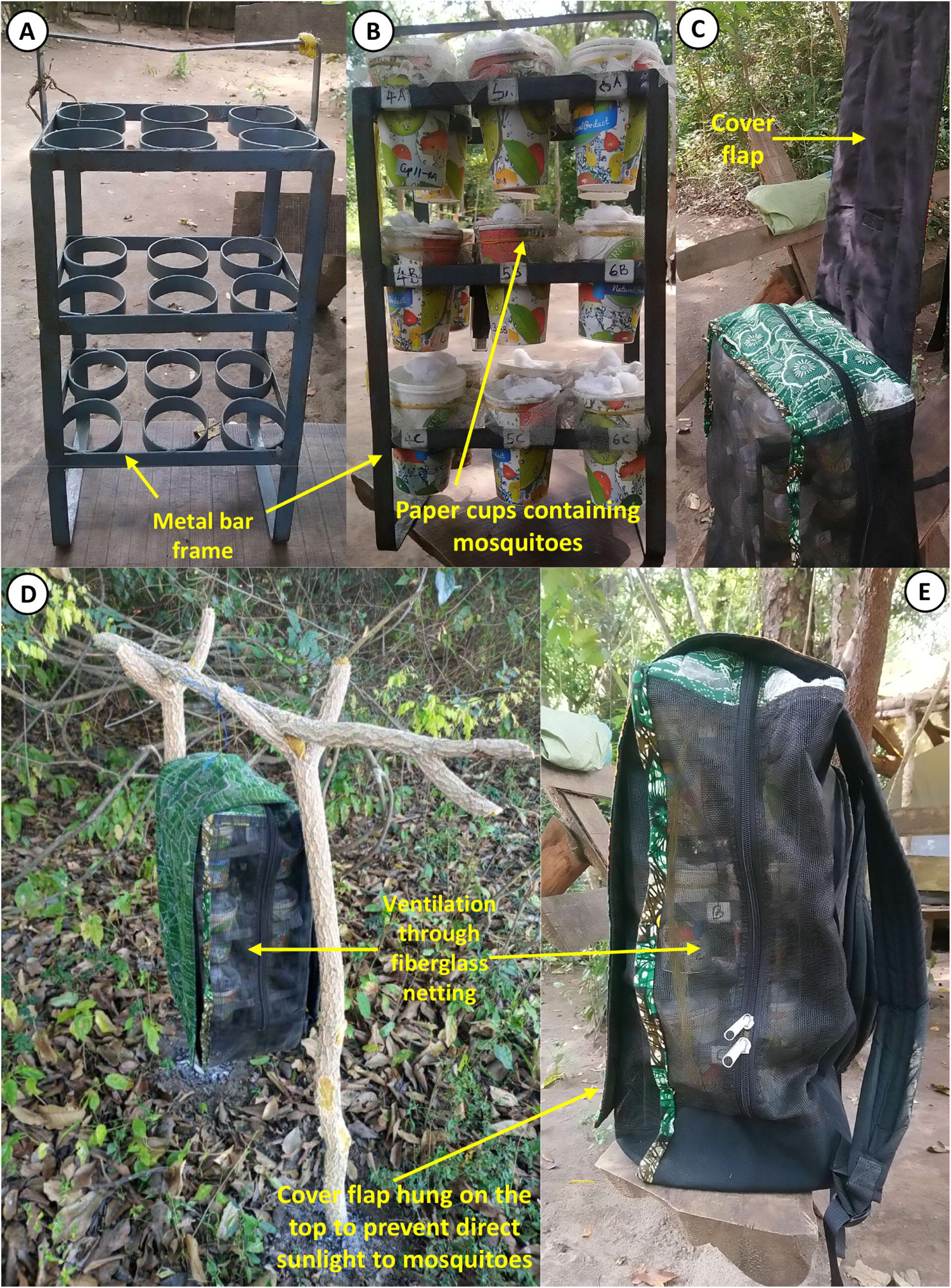
Annotated photographs of (**A**) the metal frame designed to hold papers cup cages of adult mosquitoes, (**B**) the same metal frame filled with paper cups of mosquitoes, (**C**) the metal frame full of cups placed inside the bespoke carrier backpack designed and evaluated herein, just as the wettable flap cover, comprised of absorbent cotton towelling enclosed within an envelope of *kitenge* cotton fabric, is dropped over the top and back of the backpack, and (**D**) the same backpack after the wettable flap has been dropped into place to cover the top and back of the backpack, while still leaving the cups inside fully visible and ventilated through the fibreglass netting body of the backpack. **(E)** the same backpack hung in the tree to protect the mosquitoes from ground predators such as ants.

### Study setting

The large and diverse study area illustrated in Figure 3 and described in detail below was considered ideal for addressing the broader goals of the overall project this study was nested within, with several complementary conservation biology (Duggan, 2024) and vector ecology (Walsh, 2024) objectives, as it represents a geographical gradient of natural ecosystem integrity ranging from fully domesticated land uses in the west to completely pristine natural ecosystems to the east.

**Figure 3.**
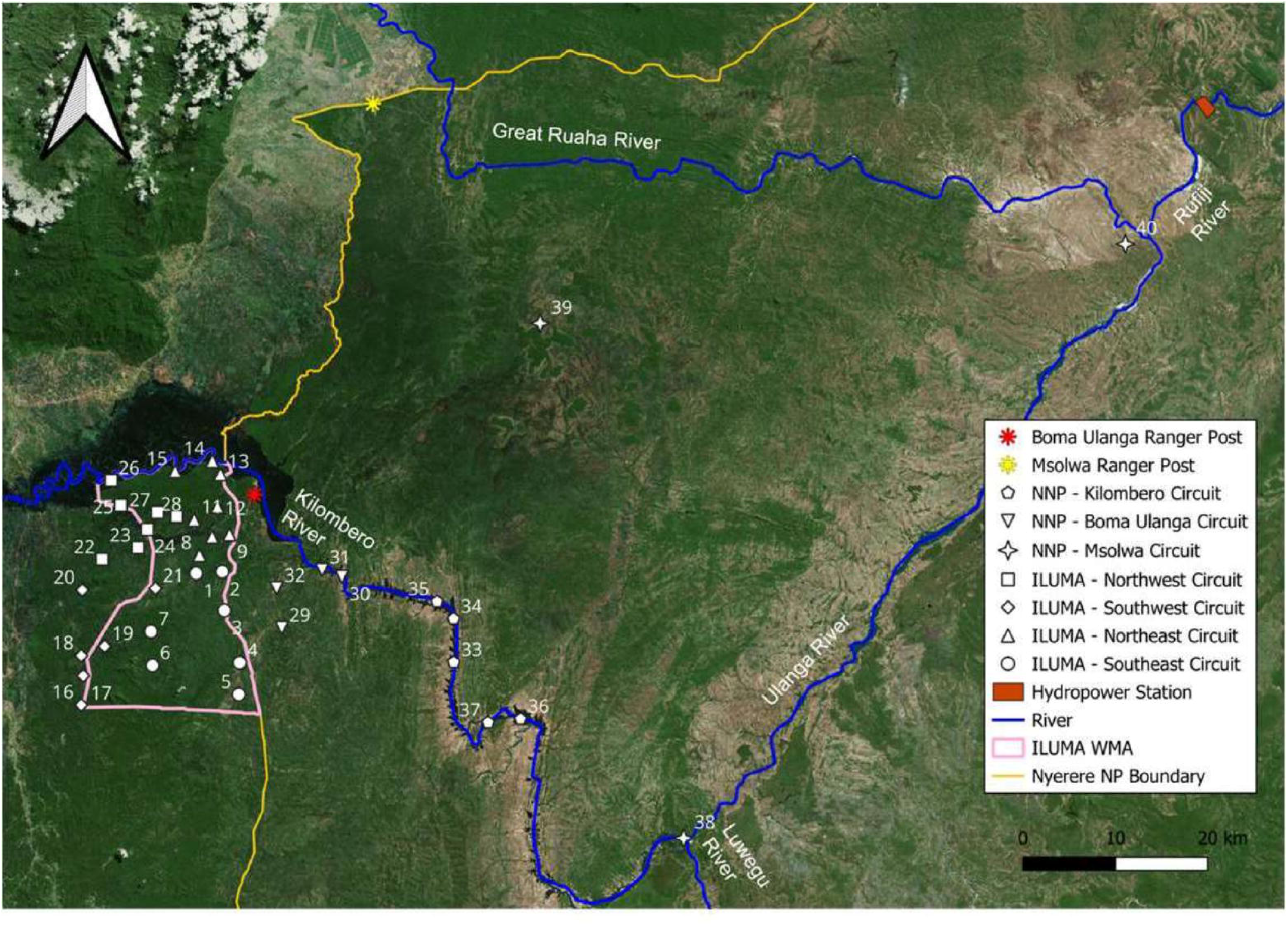
Map displaying the distribution of suitable camping locations used as the sampling frame for all the surveys described in this report. Each of the 40 camps detailed in supplementary file 1 are illustrated in the geographic context of the boundaries of the Ifakara-Lupiro-Mang’ula Wildlife Management Area (ILUMA WMA) and Nyerere National Park (NNP).

**FIGURE 4.**
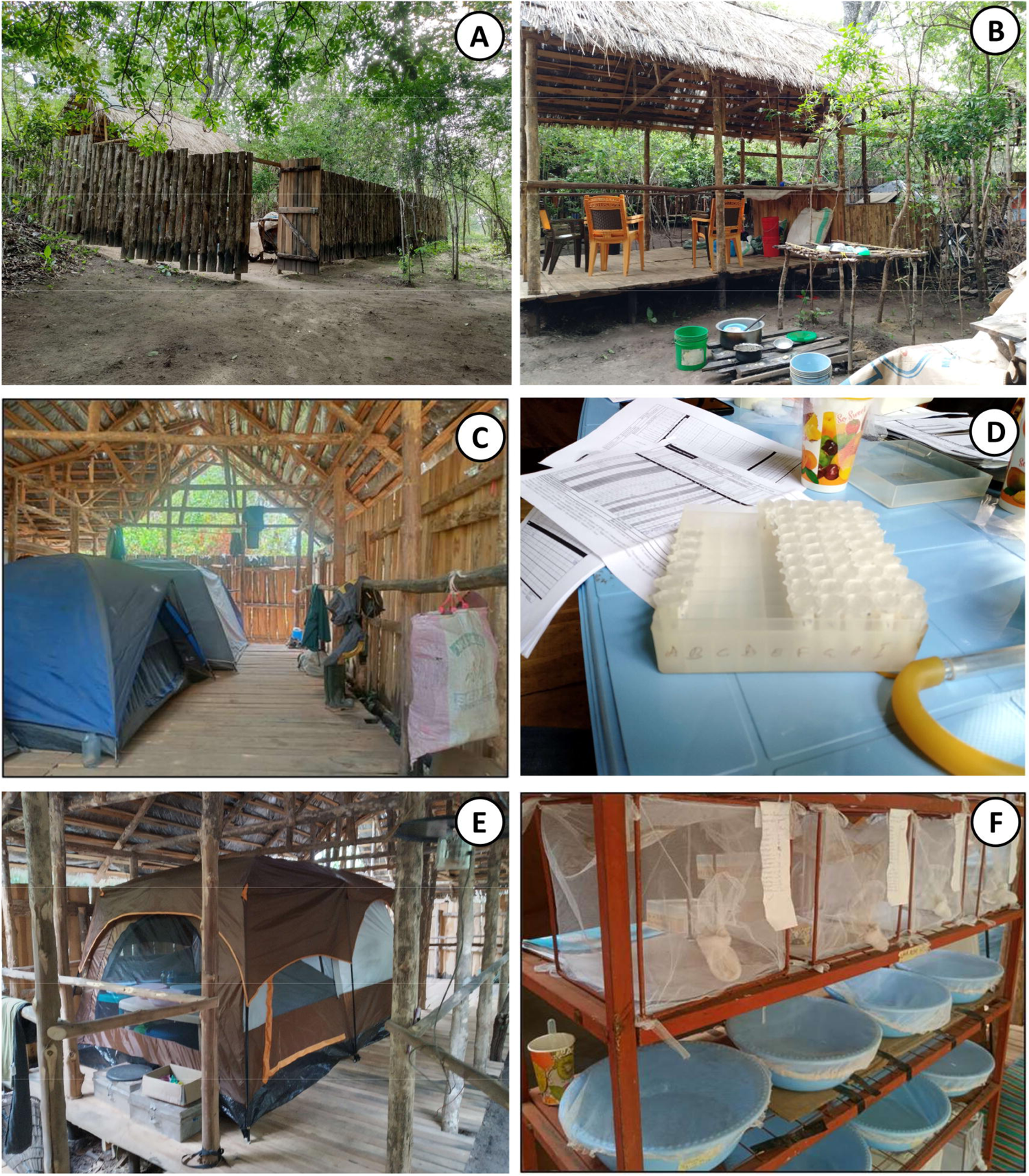
Illustrative photographs of the central camp established for the project at *Msakamba* (Figure 3). **A**: The surrounding fence for security, **B**: The kitchen and food storage area, **C**: Tents for sleeping in, **D**: An area for sorting of wild-caught mosquitoes, **E**: A field insectary tent for housing mosquito adults and larvae, **F**; Mesh cages and plastic water basins for respectively rearing adult and larvae in the field insectary.

This study was largely conducted at the downstream fringe of the inland delta of the Kilombero River (Figure 1) which is the largest low altitude wetland in east Africa, located in the Morogoro region of southern Tanzania (Mombo et al., 2011; RIS, 2002). The Kilombero valley is part of the Rufiji River Catchment Basin and is delimited by the Udzungwa mountains to the north and the Mahenge mountains to the south (RIS, 2002). The region has what are referred to as a short and a long rainy season, from approximately November to February and from March to April, respectively. The short rains are typically intermittent, sporadic and unreliable, whereas the long rains are usually more predictable and consistent. The dry season generally lasts from June to December, during which time the area experiences little to no rainfall. The malaria transmission system and vector populations of the Kilombero valley have been exceptionally well-characterised (Drakeley et al., 2000; Killeen et al., 2006; Msugupakulya et al., 2023; Russell et al., 2011), and following the disappearance of *An. gambiae s.s* after successful scale up of long-lasting insecticidal nets, the two main vectors of malaria in the area are now *An. funestus* Giles and *An. arabiensis* Patton (Kaindoa et al., 2017; Lwetoijera et al., 2014; Maia et al., 2016; Mapua et al., 2022). Located south-east of Ifakara town, lies more than 500km^2^ of community lands under the stewardships of 15 villages spanning both Kilombero and Ulanga districts which is collectively managed through the Ifakara-Lupiro-Mang’ula (ILUMA) WMA (Figures 1 and 3). It was formally established as a wildlife management area in 2015 for the purpose of practicing sustainable land use and promoting wildlife conservation in the buffer zone between a rigorously conserved state-protected area (formerly the Selous Game Reserve, now Nyerere National Park (NNP)) and adjacent village communities, while also creating opportunities for local communities to generate income and advance rural development (MNRT, 2022). Miombo woodland is the predominant natural land cover type across most of the ILUMA WMA, although a sizeable track of dense groundwater forest stretches along the south-bank of the Kilombero River. On the north-bank, floodplain grassland extends to the rainforest covered Udzungwa mountains. These very distinct ecosystems support sizeable populations of wild herbivores and carnivores, including iconic species like elephant, African buffalo, lion, leopard and African hunting dog. The western boundary of ILUMA WMA borders lands designated for agriculture and grazing extending as far as the main road connecting Ifakara and Mahenge (Figure 1 and 3). Due to its proximity to domesticated land and human activity, many areas within the ILUMA conservation area that are close to the western and southern boundaries are vulnerable to unauthorised exploitation by humans, and large sections of these areas have even been cleared for agriculture, grazing and settlement (Duggan, 2024). Other types of unauthorised human disturbance occurring across the designated conservation area, include timber harvesting, charcoal burning, fishing, and game meat hunting. However, as one moves further into ILUMA WMA and away from human settlement, natural ecosystems become less exposed to human disturbance and are increasingly inhabited by abundant populations of wild mammals that offer mosquito populations diverse alternatives to humans and livestock as potential blood sources. The eastern boundary of ILUMA WMA is marked by a road running in a north-south direction along its border with Nyerere National Park (NNP) (Figure 3). Established in 2019, NNP is Tanzania’s largest and newest national park. NNP was formerly a part of the Selous Game Reserve and covers an area of 30,893km^2^ of pristine wilderness (TANAPA, 2019). Presently, most tourist activities and infrastructure development occur in the northern section of the park, along the Rufiji River between Ikwiriri and Kisaki, that have been previously used for photographic tourism rather than big game hunting even when these areas were part of the Selous Game Reserve. At the time of the study, however, the western area of NNP that borders ILUMA WMA was only hosting limited numbers of game fishing tourists, so it experienced little human activity other than patrols by park rangers and occasional visits by poachers. The miombo woodlands of ILUMA WMA, extend eastward beyond the border road and well into the national park, providing an extensive habitat for the same species that are found in ILUMA WMA. Additionally, open grasslands, acacia scrub and undisturbed floodplains in NNP favour mammalian species that are not regularly seen in ILUMA WMA, such as impala and kudu.

### Establishment of a permanent camp and field insectary at the centre of the ILUMA Wildlife Management Area

Colonized *An. arabiensis* were obtained from the IHI central insectary facility in Ifakara (Figure 1) and transferred to a field insectary established at the central camp for this project at *Msakamba* (Camp 1 in figure 3 and supplementary file 1), where they were propagated as per standard procedures to provide offsprings for evaluating the mosquito carrier backpack described above. The *Msakamba* camp, which is named after the seasonal stream it was built alongside, was established as a logistical operations hub for the overall project that this study was nested within, the overall goal of which was to establish whether wild-caught *An. arabiensis* malaria vector mosquitoes collected from locations where humans and their livestock are scarce or absent may retain insecticide susceptibility traits that have been lost from mosquito populations in nearby towns and villages (Pinda et al., 2020; Urio et al., 2022). The camp was fenced and equipped with essential basic infrastructure like thatch roofs, tables, chairs, large tents, a kitchen, and solar-powered electricity supply (Figure 3).

Because most of the area within ILUMA wildlife management area (WMA) cannot be traversed by vehicle for large part of the year, the central location of *Msakamba* (Camp 1 in figure 3 and supplementary file 1) made it possible to reach it on foot from any other camp in ILUMA WMA within a day, so that live adult and larvae samples could usually be brought there within 48 hours of collection, regardless of the weather conditions. It should be noted, however, that this sometimes-required arduous all-day hikes of up to 25km, circumnavigate extensive flooded valleys. It also ensured that food supplies, as well as recharged batteries and power packs for mobile phones and field equipment could be regularly delivered to the mobile field team, who moved from camp to camp every two days, along circuits of up to 8 camps at a time that typically took about two weeks to survey (Duggan, 2024; Walsh, 2024). Village game scouts (VGS) recruited from the stakeholder communities, who are responsible for patrolling the ILUMA WMA and escorting any visitors to the conservation area, were also engaged as essential team members for this study (Duggan, 2024; Walsh, 2024). *Msakamba* was occupied and maintained on a permanent basis throughout the study, by a team of VGS and technicians, who were responsible for maintaining the field insectary, rearing field-caught mosquitoes and carrying out experimental assessments of their insecticide susceptibility phenotypes. While the original study design for the overall project included only sampling locations (referred to herein as *camps* because each survey visit involved setting up a temporary camp there for two nights) within the WMA or the neighbouring villages immediately to the west (Duggan, 2024; Walsh, 2024), it was subsequently expanded to include 12 additional camps distributed across a much larger area within Nyerere National Park (NNP) to the east (Figure 3).

### Experimental assessment of the backpack with insectary-reared mosquitoes

Ten insectary-reared female *An. arabiensis* mosquitoes were placed in each of six labelled paper cups, each with a wedding veil netting cover secured with a rubber band on top of the cup. Each cup of the mosquito used over the course of the study was labelled with a unique serial number, which matched to a data collection form with the same number for recording mortality events, escapes, and other censoring events. Mosquitoes were provided with a 10% glucose solution soaked into a ball of a cotton wool that was placed on top of the netting cover. Each time a new batch of mosquitoes from the insectary was added to the experiment, 18 pieces of paper were each labelled with one of the 18 metal frame position numbers and each of the six cups of mosquitoes was allocated to a random position in the frame by blindly picking a piece of paper from among the 18, reading the position number, and placing the paper cup in that specific position. At the end of the first randomization, six positions out of the 18 were occupied and that set of six cups, all originating from a single batch of mosquito from *Msakamba* insectary, was considered to represent one replicate unit of the experiment.

This process of randomization without replacement was repeated approximately every four days, at which point the field team completed the experimental cycle of visiting one camp other than *Msakamba,* until every position within the two metal frames for the two backpacks were collectively filled with six batches of six cups per batch, with 10 mosquitoes in each cup. At the outset, an EasyLog (500-711, EL-USB-2-LCD Lascar Electronics Ltd) temperature and humidity recorder was also placed inside the carrier backpack, and it was moved through various positions within the backpack, at the top, middle, and bottom of the metal frame, every experimental day. Another logger was hung immediately outside the backpack to record temperature and humidity just outside it. For each experimental replicate, with a new batch of mosquitoes from the insectary, three additional cups containing 10 mosquitoes each were treated identically except they were kept in the field insectary and maintained there under normal rearing conditions, to serve as a control group for each batch that was not transported on foot across the study area.

### Transportation and mortality monitoring of insectary reared mosquitoes

Nine different locations (Table 1) were chosen within the ILUMA WMA, as representative destinations that the field team transported different batches of insectary reared mosquitoes on foot to and from using the mosquito carrier backpack. The field team spent approximately four nights at each of the mobile camps visited during which time they carried out extensive hikes around the surrounding areas while carrying the backpack of mosquitoes with them. During periods spent each day in a specific camp, or while resting in the afternoons during hikes around the area surrounding the camp, the mosquito backpack was hung off the ground under the shade of a tree, on the end of a piece of string that was tied to a suitable branch tree, to keep the samples off the ground and avoid predation by ants (figure 2D). Note also that the backpack was kept at a minimum distance of 20m from the campfire, to avoid mortality caused by smoke. The absorbent top cover flap was soaked with water at least each morning and evening to ensure optimal environmental condition for the mosquitoes inside the backpack. The field team recorded number of dead mosquitoes, and any censoring event like escapes or removal by ants, in each cup every morning and evening. These recordings continued until no living adults remain in the cup or the study was terminated, when the last batch of mosquitoes had been transported around the WMA for ten days. The same mortality monitoring process was applied to the control cups that were left in the field insectary at *Msakamba* and maintained under standard rearing conditions.

**Table 1:** Camp names and numbers of camps visited within ILUMA during the experimental transportation of insectary-reared mosquitoes around the WMA in the carrier backpack illustrated in figure 2, so that their survival under such conditions could be assessed. The approximated distance from the central camp *Msakamba* to each of the camps listed below, and total distance travelled by each batch (1 to 6) of mosquito are also indicated.

| Camp number<sup>a</sup> | Camp name | Distance from *Msakamba* main camp (km) | Total distance travelled for each batch (km) | Mosquito batch number |
| --- | --- | --- | --- | --- |
| 2 | Bwawa la msiba wa Deo | 3.1 | 143 | 1 |
| 10 | Bwawa la Miembemi | 7.8 | 137 | 2 |
| 12 | Bwawa la Maya | 7.9 | 121 | 3 |
| 28 | Bwawa la Semka | 6.5 | 106 | 4 |
| 14 | Mikeregembe | 12.3 | 93 | 5 |
| 7 | Bwawa la Chakacheni | 8.0 | 68 | 6 |
| 4 | Bwawa la Nandete | 10.0 | 68<sup>b</sup> | 6 |
| 3 | Bwawa la Nyati | 5.0 | 68<sup>b</sup> | 6 |
| 26 | Bwawa la Mamba Luhogi | 11.0 | 68<sup>b</sup> | 6 |

<sup>a</sup> See supplementary file 1.
<sup>b</sup> Batch 6 was the last batch of insectary mosquitoes transported from *Msakamba* and it went through the rest of the camps (3, 4, and 26) together with all other batches.

After four or more days, the field team returned to the main camp and rested for the night. The following morning, another batch of six paper cups, all labelled with their own new consecutive serial numbers, was added to the backpack by following the same procedures and randomization process described above. The field team then departed again to the next mobile camp and spent four nights recording mortality in each cup every morning and evening on either side of extended daily hikes.

These cycles were repeated until six replicates of batches of six cups with 10 mosquitoes each were transported over a cumulative distance ranging from 68 to 143 km over 10 to 25 days. Also, the corresponding three replicates of 3 paper cups of mosquitoes were left in the field insectary as a control group for each mosquito batch or experimental replicate for a corresponding duration. Therefore, the whole experiment consisted of 36 transported cups of mosquitoes and 9 control cups, collectively spanning six different batches of mosquitoes from the field insectary.

### Practical application of the carrier backpack to maintenance and transportation of field caught mosquitoes

The overall project that the backpack described herein was designed to enable was carried out using a rolling cross-sectional design, with four rounds of surveys encompassing a total of 40 defined camp locations across NNP, ILUMA WMA and the villages immediately to the west of it over the course of two years (Figure 3 and supplementary file 1). Most sampling was carried out during the wet season as rainfall generally increases *Anopheles* mosquito abundance across most locations (Gillies & De Meillon, 1969) including this one (Charlwood et al., 1995; Gillies & De Meillon, 1968). Adult mosquito surveys, together with complimentary larval surveys (Walsh, 2024), land use and wildlife activities (Duggan, 2024), were completed sequentially at the 40 defined camp locations that encompassed a range of land uses and ecosystem integrity states with a wide range of mammalian species abundance and diversity (Duggan, 2024; Walsh, 2024) (Figure 3, supplementary file 1).

For the original protocol, a total of 28 camps were selected after scouting potential locations in ILUMA WMA and considering the suggestions of the VGS based on their vast personal knowledge of the area. The broad geographic distribution of the camps was planned to encompass as wide of a range of ecosystem states as possible, by including all parts of ILUMA WMA and the neighbouring domesticated land to the west. However, the exact position of a camp location was ultimately determined by the requirement for perennial surface water that *Anopheles* larvae and adults could be collected from. Crucially, the availability of water for cooking, drinking, and bathing was essential for sustaining the field team in the absence of regular vehicle support. In most cases, the selection of camp locations with accessible surface water also ensured that firewood and at least some shades were available in situ. Accessibility by foot during the wet season was also a critical factor to consider, to ensure that each camp could be safely reached, and the transport of live mosquitoes could be completed even during periods of heavy rain and flooding. The presence of one or more glades or valleys with numerous perennial waterbodies, like waterholes, ponds and streambeds near the camp was also required so that *Anopheles* mosquito larvae (Walsh, 2024) and adults.

Once the selection process was complete, each camp was given a unique number and a name and the full list of camps was then divided into quarterly *circuits* that roughly corresponded to the *southeast*, *southwest*, *northeast*, and *northwest* of the ILUMA WMA, with each circuit named accordingly (Figure 3, supplementary file 1). The circuits were designed to manage the physical challenges of the study, and so subsets of 6 to 8 camps were grouped into distinct circuits based on their geographic proximity to one another and the practicality of visiting them all in a circular route that started and ended centrally at the *Msakamba* camp. Each circuit had a planned route that minimised walking distances between camps, to ensure that the team had enough time to rest in the afternoon before proceeding with data collection later into the evenings and in the early mornings. Camps were visited consecutively for two nights each and involved intensive daily data collection procedures. Therefore, the circuit design for sequentially visiting camps in a rolling cross-sectional survey was crucial for sustaining optimal long-term data collection by limiting investigator fatigue and allowing them to take breaks of a few days in between periods of continuous field work that typically lasted about two weeks for a single circuit.

The original protocol planned for a total of three rounds to be completed from January to March, March to May, and August to October 2022, representing the short rainy season, long rainy season, and dry season, respectively. It was planned that each round of surveys in the longitudinal rolling cross-sectional study design would visit all 28 camps detailed in supplementary table 1, but a few camps were omitted from each round for pragmatic reasons, such as a lack of surface water during the dry season or inaccessibility due to severe flooding during intense rains.

However, no camp in the ILUMA WMA appeared to lack signs of human disturbance, and only a few remained relatively pristine, so four new mobile camps inside NNP were added (number 29-32, figure 3) to the study design at the end of round three in November 2022, forming a new circuit that was surveyed with vehicle support for logistical and safety reasons. These camps were located inside the boundary of NNP, immediately to the east of ILUMA WMA to capture absolute pristine environments and were accessed by vehicle via the NNP ranger post at *Boma Ulanga*, for which that circuit was named. These were then repeated at the start of the fourth and final round of data collection that was completed from February to July 2023, representing the whole wet season and the beginning of the dry season for that calendar year. Considering the initial unexpected sibling species composition and insecticide phenotype results obtained from the first four NNP camps, it was decided to extend the sampling frame deeper into the park and adjust the field protocol to collect and immediately preserve additional collections of larvae that would not be used to rear adults in the *Msakamba* insectary (Walsh, 2024).

To obtain samples as far away from human beings, and as deep into pristine ecosystems as possible, another, eight camps inside NNP were added to the end of the fourth and final round. First, an additional subset of five camps that were accessed by boat via the Kilombero river, and correspondingly named the *Kilombero circuit,* were visited (Figure 3). Pushing even further east into the park, camps 38 to 40 that were accessed by vehicle along main park roads via the Msolwa ranger post and were therefore named accordingly as the *Msolwa circuit* (Figure 3).

Live mosquitoes were trapped using two methods: a netting barrier interception screen (Figure 6) similar to that developed for sampling exophilic mosquitoes in the Pacific (Burkot et al., 2013) and Centres of Disease and Control (CDC) light traps (John W. Hock Company, product number 512). After arriving at a new camp location, a shaded area next to a water source was scouted to identify a suitable place for setting up the tents, cooking, resting and processing mosquitoes. The batteries for four CDC light traps that were used for trapping live mosquitoes were then charged as soon as possible using portable solar panels. On overcast days, especially during the rainy season, charged batteries were received from the *Msakamba* central camp every two days by the team responsible for taking the collected mosquito samples. A site within a valley or open natural glade, at an approximate distance of 100m from the camp, was identified and a five-panel 25m long interception screen made from standard mosquito netting was set up as shown in figure 5.

**FIGURE 5.**
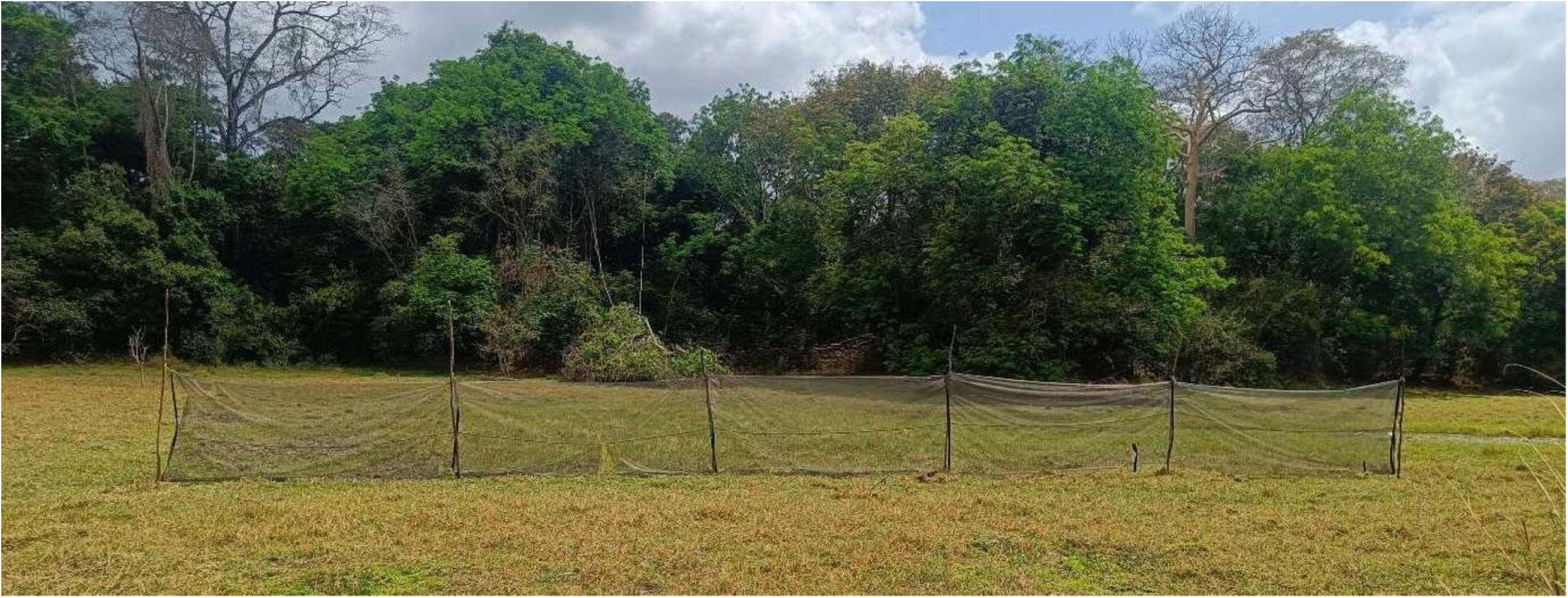
A photograph illustrating a typical set up for the screen netting interception screen trap in a typical open natural glade surrounded by dense woodland.

**FIGURE 6.**
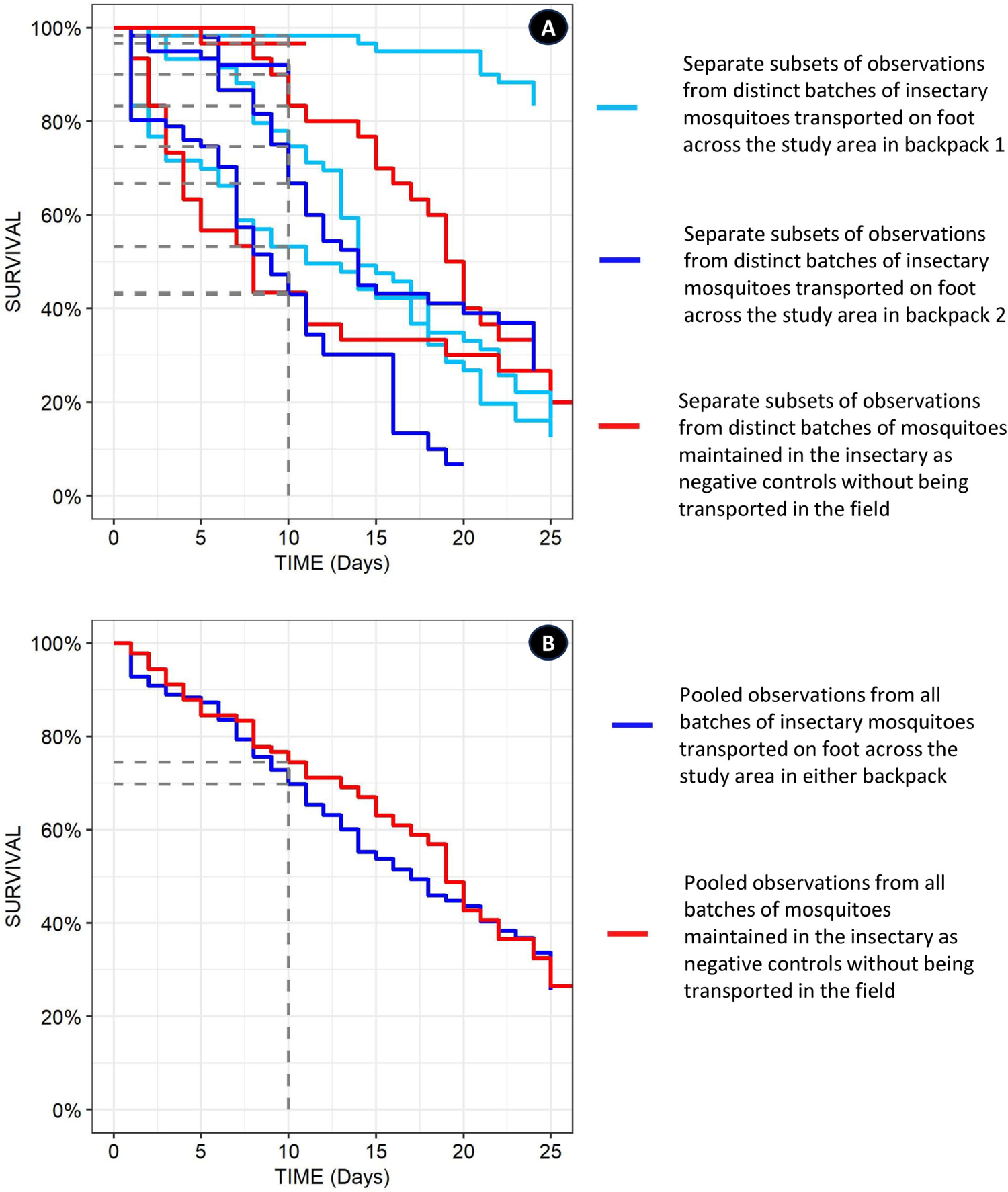
Observed survival curves from the experimentally controlled assessment, in which insectary-reared *Anopheles arabiensis* mosquitoes were maintained inside carrier backpacks (Figure 2) over extended periods of transport on foot, compared with negative controls that were not transported but rather kept in the insectary (Figure 4D, E and F). Panel **A** illustrates the considerable variance between and covariance within batches of mosquitoes assigned to either long distance transport on foot inside the backpack or standard maintenance inside the insectary. Panel **B** illustrates how the overall mean survival curves derived from the pooled data for all the different cups and mosquito batches in the backpacks do not differ from those of the negative controls that were kept in the insectary (P=0.16).

Collections at the interception screen were carried out five times after dusk for every hour between 19:30 and 23:30, and once just before dawn at 05:30, specifically to target the known peak outdoor biting activity of adult *An. arabiensis* in the Kilombero valley (Maia et al., 2016; Russell et al., 2011). Both sides of the screen were scanned in succession, using a torch to carefully inspect each panel in an up-and-down motion for resting mosquitoes. Once identified, the mosquito was suctioned into a collection cup using a custom-made, standard *Prokopack* aspirator, which is a handheld aspirator powered with a 6V GWESPECS battery (Maia et al., 2011; Vazquez-Prokopec et al., 2009). Separate collection cups were used for every hour and were immediately labelled with the date and time of sampling. A small ball of cotton wool was soaked into a 10% w/v glucose solution and placed on top of the collection cup as a food source.

Additionally, four CDC light traps ran overnight between 19:00 and 07:00, each in a different location to capture any potential variability in host-seeking behaviour or dispersal patterns.

Each trap was set up in one of four distinct ways: One suspended in an open valley or natural glade, one next to a stream bed, one in the camp site, and one next to an occupied tent. Again, the traps inside the camp and next to the tent were placed as far away from the campfire as possible. The following morning at 07:00, all captured mosquitoes from the interception screen and the CDC light traps were transferred into clean, labelled paper cups using a mouth aspirator, supplied with fresh glucose solution soaked into cotton wool.

Then an experimental design form template (ED1) from a standardised mosquito collection informatics platform (Kiware et al., 2016) was completed to include variables such as *date*, *camp number*, *collection method*, *habitat type* (referring to where the light trap or barrier screen was placed)*, start time*, *finish time*, *experiment day*, *volunteer initials* and any further *comments*. Samples in the paper cups were labelled with the date and time, as well as the ED1 form *serial number*, *form row* and habitat type values, for sample processing and tracing in the field insectary at *Msakamba.* These cups containing mosquitoes were then placed back in the mosquito carrier backpack (Figure 2), following which the cover flap was soaked in water exactly as described earlier for the experimental assessment using insectary reared colony mosquitoes.

After the mosquitoes were collected and prepared on the first morning, this procedure was repeated at the same camp the following evening and morning after recharging the batteries in the afternoon. On this second morning, the field team would break camp and hike with all the equipment to the next camp location to be surveyed, but only after two VGS arriving from the *Msakamba* central camp delivered supplies and collected the backpack of mosquito samples (figure 2E) to return them to the field insectary for sorting, counting, and rearing. After arriving at the new mobile camp location, the routine described above was repeated until the relevant survey circuit was competed.

### Statistical analysis

All statistical analyses were performed by using the *R*^®^ version 4.1.3 open-source software package, through the *Rstudio*^®^, version 2023.09.1.494 environment (Team, 2022). To estimate survival of insectary-reared female *An. arabiensis* mosquitoes, Cox regression mixed effect models were fitted to the data with the *coxme* package, treating mosquito mortality, or alternatively censoring events like escapes, predation by ants or termination of the study as the dependent response variable, while treatment (transported or non-transported) was included as the independent variable of primary interest, whereas cage position, and cup identifiers nested within insectary mosquito batch were treated as crossed random effects.

Furthermore, to compare survival rates across different rounds of field collections, a logistic GLMM was fitted to the relevant observations of mosquitoes collected in the field and then transported to the *Msakamba* insectary, using the *glmer* function of the *lme4* package, treating mosquito survival as the binomial response variable, round of collection and area (ILUMA versus NNP) as categorical fixed effect predictors, and camp number as a random effect. However, to estimate the mean survival rates for wild mosquitoes obtained with different methods of collection, an otherwise identical GLMM model but lacking an intercept was fitted to the same data, with survival as the binomial response variable, while method of collection and camp number were respectively treated as the sole categorical predictor and random effect variables.

## RESULTS

### Experimental assessments using insectary reared *Anopheles arabiensis*

Mixed effects Cox regression, allowing for considerable covariance and variance between different batches of *An. arabiensis* colony mosquitoes from the insectary, which corresponded to quite distinctive experimental replicates in terms of their survival curves (Figure 6A) indicates that transporting mosquitoes in the backpack up to 25 days made no consistent difference (P=0.16) to their overall longevity (Figure 6B). On average, across both carrier backpacks combined, at least 70% of insectary mosquitoes were still alive after 10 days of transportation. Such high survival rates over such extended periods went far beyond the original ambition of the study, which was to keep the majority of mosquitoes alive for at least 3 days in the field. While different levels and positions within the backpack had no appreciable effect upon mosquito survival (P=0.20 and 0.16, respectively, for the contrasts in goodness of fit between alternative models that did and did not include these variables as random effects), considerable variation occurred between individual cups of mosquitoes (σ = 0.5953, P= 0.71 for the improvement in goodness of fit achieved by including this random effect) and between different batches of mosquitoes from the insectary (σ = 1.3798, P= 0.16 for the improvement in goodness of fit achieved by including this random effect).

Inside the backpacks, the temperature and relative humidity were maintained approximately within the standard recommended ranges of 17°C to 27°C and 80 ± 10% respectively, for controlled rearing conditions in insectaries. Therefore, the original intention to develop a self-cooling and self-humidified mosquito carrier backpack was apparently achieved. Indeed, conditions inside the backpack while in use over long journeys, covering up to 143km over up to 25 days, were generally very similar to those in the insectary (Figure 7). In fact, daily maximum temperatures inside the backpack were generally milder than in the insectary (figure 7A) while humidity levels were usually the same or higher (Figure 7B). Perhaps unsurprisingly, environmental conditions inside the backpack and in the insectary were generally less harsh than in the external environment immediately outside the backpack (Figure 7).

**FIGURE 7.**
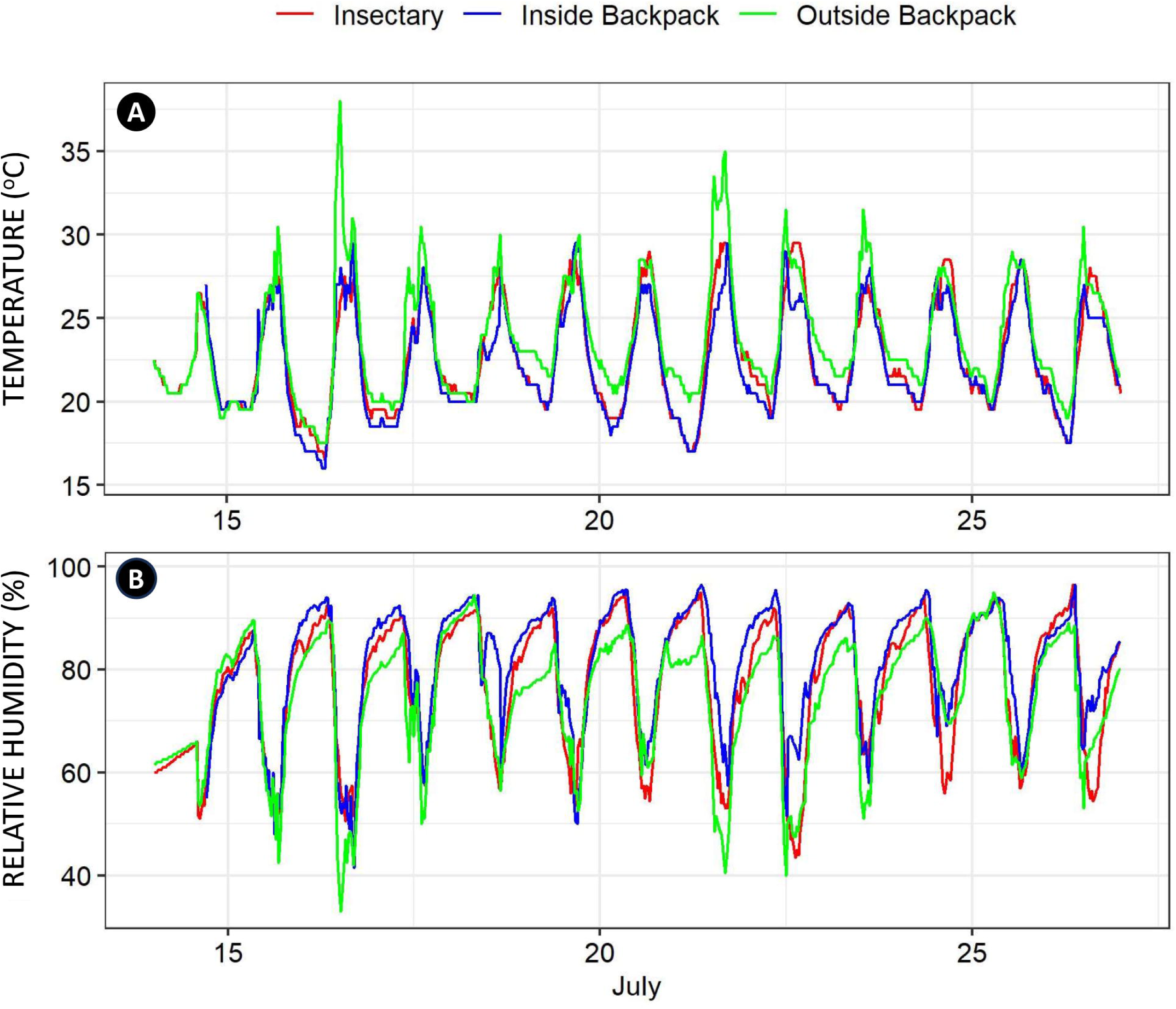
Variations in temperature (panel **A**) and humidity (panel **B**) recorded every 30 minutes by three data loggers over the course of 25 days and approximately 143km of mosquito transport on foot across the study site (Table 1 and Figure 3), with one device placed inside the mosquito carrier backpack, another immediately outside mosquito carrier backpack and another inside the insectary where the negative control mosquitoes were kept over the same period.

### Practical application of the carrier backpack to maintenance and transportation of field caught mosquitoes

The overall mean [and 95% confidence interval] survival of wild-collected *Anopheles gambiae* complex mosquitoes, from all the scattered points of collection across the study site back to *Msakamba* (Figure 4), estimated by fitting a GLMM was 58.2% [47.5, 68.2%]. Survival was somewhat higher (P=0.0039) by GLMM in round three than in any of the other survey rounds (Figure 8A). This was probably because most of the troubleshooting and optimization of the field procedures for collecting mosquitoes and maintaining them in the carrier backpack had been completed by round three and the scientific team in the mobile field team remained at full strength up to that point: While there were three professional postgraduate scientists accompanying the VGS in the field throughout the first three rounds of surveys, so that the supervision of their work was distributed shared to avoid fatigue and ensure consistency, in the fourth round this was reduced to only one. Surprisingly, no difference in survival (P=0.74 by GLMM) was observed between mosquitoes collected in ILUMA WMA and those collected in NNP (Figure 8B), even though transport back from NNP involved travel over much longer distances (Figure 3), sometimes involving lengthy journeys by car or boat. Overall, mosquitoes collected from the barrier screen collection had a higher mean survival rate than those collected with CDC light traps (64.5% [58.2, 70.5%) versus 50.0% [43.5,56.5%], respectively, P<<0.00001 by GLMM analysis) figure 8C.

**FIGURE 8.**
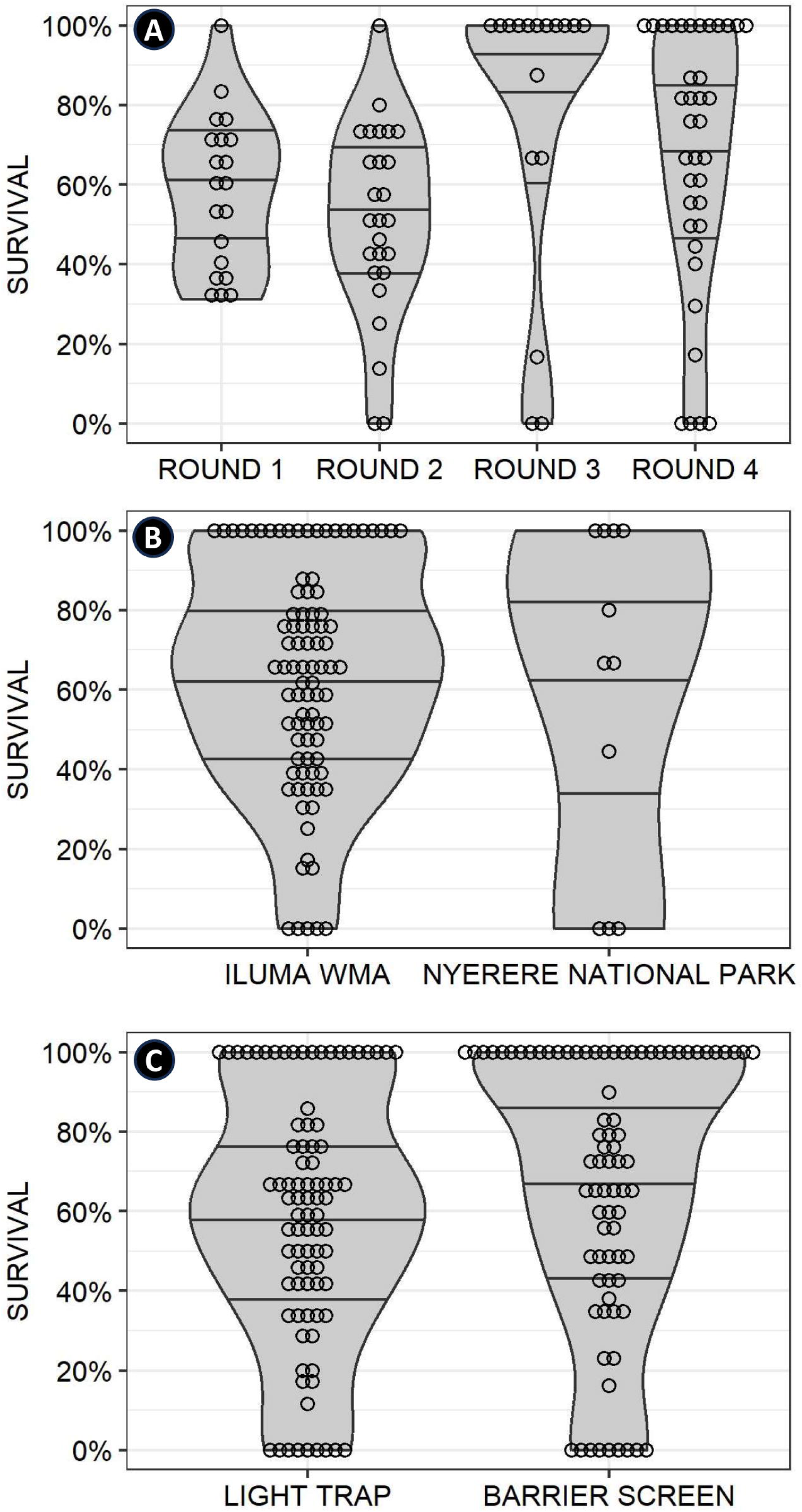
Dot plots of the distributions, medians and interquartile ranges for the observed survival rates for each two-night collection of wild mosquitoes at each camp during each of four distinct survey rounds, over the period between being captured in the field and then maintained and transported back to the field insectary at *Msakamba* (Figure 4) from all across the study site (Figure 3) within the mosquito carrier backpack (Figure 2). **A**: Comparing the four distinct rounds of surveys and associated mosquito collections. **B**: Comparing mosquito collections made in or immediately to the west of the Ifakara-Lupiro-Mang’ula Wildlife Management Area (ILUMA WMA) and Nyerere National Park (NNP). **C**: Comparing the two different mosquito capture methods used.

## Discussion

This study clearly demonstrates that it is possible to transport female *An. arabiensis* mosquitoes up to 143km on foot under challenging wilderness field conditions across a 509 km^2^ WMA for up to 25 days using a mosquito carrier backpack, with a 70% survival rate sustained consistently over ten days. When put into practical use in a much bigger study, spanning an even larger (>4000km^2^) and more diverse survey area (Walsh, 2024), the majority of mosquitoes caught were successfully returned alive to the central insectary, for further investigation of their heritable insecticide susceptibility traits. This mosquito carrier backpack provided mosquitoes with favourable environmental conditions that compared well without our permanent field insectary, and it is an important advantage that it can readily made locally with affordable materials that are widely available across rural Africa. One complete unit of the mosquito carrier backpack costs a maximum of $70 to fabricate and weighs no more than 5 kg when it is fully filled with paper cups and mosquitoes, making it quite affordable, lightweight, and convenient for practical use.

Beyond addressing our project’s specific needs, the backpack described herein may also have other uses. For example, the implementation of the sterile insect technique (SIT) for mosquito control always involve the transportation of live specimens from the production facility to the release points and sometimes beyond the reach of motorized transport (Ernawan et al., 2022; Gómez et al., 2022). Specifically, for SIT procedures, other factors apart from transportation such as packaging and storage conditions may affect the survival and longevity of mosquitoes(Chung et al., 2018; Sasmita et al., 2022), and any potential role for carrier backpacks based on this prototype would probably be limited to parts of remote rural areas or dense urban settlements that may be more readily accessed on foot. While transportation of sterile male *An. arabiensis* under low temperature did not significantly affect their survival (Culbert et al., 2017) and mating competitiveness (Helinski et al., 2008) over a period of up to 24 hours, it may not always be access or maintain refrigeration across extensive, often remote rural areas.

To the authors’ knowledge, this is possibly the only study that succeeded in transporting insectary and wild-caught female *An. arabiensis* on foot over such long distances under such challenging field conditions, and it is encouraging that such high survival rates were achieved. Nevertheless, further experiments should be conducted to assess how well other important phenotypic characteristics, such as feeding rate, mating competitiveness, and oviposition rate are maintained after long-distance transportation on foot using this new carrier backpack prototype.

## AUTHOR CONTRIBUTIONS

Deogratius R. Kavishe: Conceptualization, investigation, methodology, formal analysis, writing original draft of the manuscript. Rogath V. Msoffe: Conceptualization, investigation, methodology, implementation review and editing. Goodluck Z. Malika, Katrina A. Walsh, Lily M. Duggan and Lucia J. Tarimo: Implementation, review, editing, validation. Fidelma Butler: review, editing, validation. Emmanuel Kaindoa: review, editing and validation. Halfan S. Ngowo: Initial statistical analyses, review editing, validation. Gerry F. Killeen: Acquisition of fund, Conceptualization, investigation, visualization, methodology, formal analysis, review, editing and validation. All authors read and approved the final submitted version of the manuscript.

## ACKNOWLEDGEMENTS

The authors wish to thank the Village Game Scouts of ILUMA WMA, for their hard work and participation in field activities. We also thank all the governance, management, and stakeholder communities of the ILUMA WMA for all their collaboration and kind assistance over the course of the study. Furthermore, we thank Prof Nico Govella, Mr Frederic Masanja, Prof Honorati Masanja, Ms Catherine Ringo, Mr Fadhili Songo, Ms Elaine Kelly, Dr Ronan Hennessy, Ms Leonie O’Doherty, Ms Sonia Montero and Prof Sarah Culloty for all the essential institutional support provided by the Ifakara Health Institute and University College Cork over the course of the study. A very special word of thanks is due to our recently deceased friend and colleague, Mr Octavian Malopola, without whom this work could never have begun, much less completed safely and successfully.

## DATA AVAILABILITY STATEMENT

**Supplementary file 1;** Table S1: The number, name, location, and ecological characteristics of each camp location, together with the circuit to which it was assigned and the number of times it was surveyed https://zenodo.org/doi/10.5281/zenodo.10946752

**Supplementary File 2**; The full data set used for the analysis presented herein https://zenodo.org/doi/10.5281/zenodo.10939877

