## Supplementary table 1 for "A self-cooling self-humidifying mosquito carrier backpack for transporting live adult mosquitoes on foot over long distances under challenging field conditions"

**SUPPLEMENTARY TABLE S1** Each camp number and name with the corresponding circuit, location and the number of rounds they were surveyed in. See [WALSH ET AL] for further details.

| *Number* | *Camp name* | *Circuit* | *Location* | *Historical Landcover* | *Round* |
| --- | --- | --- | --- | --- | --- |
| 1 | Msakamba | Southeast | Inside ILUMA | Miombo woodland | 1,2,3,4 |
| 2 | Bwawa la Msiba wa Deo | Southeast | Inside ILUMA | Miombo woodland | 1,2,3,4 |
| 3 | Bwawa la Nyati | Southeast | Inside ILUMA | Miombo woodland | 1,2,3,4 |
| 4 | Bwawa la Nandete | Southeast | Inside ILUMA | Miombo woodland | 1,2,3,4 |
| 5 | Korongo la Bundu | Southeast | Inside ILUMA | Miombo woodland | 1,2,3,4 |
| 6 | Bwawa la Namamba | Southeast | Inside ILUMA | Miombo woodland | 1,2,3,4 |
| 7 | Bwawa la Chakacheni | Southeast | Inside ILUMA | Miombo woodland | 1,2,3,4 |
| 8 | Bwawa la Njuju | Northeast | Inside ILUMA | Miombo woodland | 1,2,3,4 |
| 9 | Bwawa la Chamvi | Northeast | Inside ILUMA | Miombo woodland | 1,2,3,4 |
| 10 | Bwawa la Miembeni | Northeast | Inside ILUMA | Miombo woodland | 2,3,4 |
| 11 | Kisima cha Seba | Northeast | Inside ILUMA | Miombo woodland | 1,2,4 |
| 12 | Bwawa la Maya | Northeast | Inside ILUMA | Miombo woodland | 1,2,3,4 |
| 13 | Bwawa la Mrope | Northeast | Inside ILUMA | Groundwater forest | 1,2,4 |
| 14 | Mikeregembe | Northeast | Inside ILUMA | Groundwater forest | 1,2,3,4 |
| 15 | Mdalangwila | Northeast | Inside ILUMA | Groundwater forest | 1,2,3,4 |
| 16 | Bwawa la Mnyuamachi | Southwest | Outside ILUMA | Miombo woodland | 1,2,3,4 |
| 17 | Tuliza Moyo | Southwest | Outside ILUMA | Miombo woodland | 1,2,3,4 |
| 18 | Mavimba Porini | Southwest | Outside ILUMA | Miombo woodland | 1,2,3,4 |
| 19 | Bwawa la Selesusi | Southwest | Inside ILUMA | Miombo woodland | 1,2,3,4 |
| 20 | Makingi | Southwest | Outside ILUMA | Miombo woodland | 1,2,3,4 |
| 21 | Bwawa la Mpunga | Southwest | Inside ILUMA | Miombo woodland | 1,2,3,4 |
| 22 | Kisaki | Northwest | Outside ILUMA | Miombo woodland | 2,3,4 |
| 23 | Uwanja wa Ndege | Northwest | Outside ILUMA | Miombo woodland | 2,3,4 |
| 24 | Bwawa la Mkwajuni | Northwest | Inside ILUMA | Miombo woodland | 2,3,4 |
| 25 | Bwawa la Mamba Luhogi | Northwest | Inside ILUMA | Miombo woodland | 2,3,4 |
| 26 | Funga | Northwest | Inside ILUMA | Groundwater forest | 2,3,4 |
| 27 | Bwawa la Mlenda | Northwest | Inside ILUMA | Miombo woodland | 2,3,4 |
| 28 | Bwawa la Semka | Northwest | Inside ILUMA | Groundwater forest | 2,3,4 |
| 29 | Kambi ya Simba | Boma Ulanga | Inside NNP | Miombo woodland | 3,4 |
| 30 | Bwawa la Kiboko Zanzibar | Boma Ulanga | Inside NNP | Acacia savanna | 3,4 |
| 31 | Zanzibar | Boma Ulanga | Inside NNP | Miombo woodland | 3,4 |
| 32 | Bwawa la Moto | Boma Ulanga | Inside NNP | Miombo woodland | 3,4 |
| 33 | Kambi ya Mamba | Kilombero | Inside NNP | Acacia savanna | 4 |
| 34 | Kambi ya Machuma | Kilombero | Inside NNP | Acacia savanna | 4 |
| 35 | Serengeti Ndogo | Kilombero | Inside NNP | Acacia savanna | 4 |
| 36 | Kambi ya Makutano | Kilombero | Inside NNP | Acacia savanna | 4 |
| 37 | Kambi ya Mawe | Kilombero | Inside NNP | Acacia savanna | 4 |
| 38 | Shughuli kubwa | Msolwa | Inside NNP | Miombo woodland | 4 |
| 39 | Bwawa la Chatu | Msolwa | Inside NNP | Miombo woodland | 4 |
| 40 | Bwawa la Umeme | Msolwa | Inside NNP | Acacia savanna | 4 |

ILUMA: The conservation area of the Ifakara-Lupiro-Mangula Wildlife Management Area

NNP: Nyerere National Park
